# Segmentation in tapeworms as a modified form of flatworm posterior regeneration involving Wnt and Hedgehog signalling

**DOI:** 10.1101/2023.10.02.560491

**Authors:** Francesca Jarero, Andrew Baillie, Nick Riddiford, Jimena Montagne, Uriel Koziol, Peter D. Olson

**Affiliations:** Department of Life Sciences, Natural History Museum, London SW7 5BD, United Kingdom; Department of Genetics, Evolution and Environment, University College London WC1E 6BT, United Kingdom; Sección Biología Celular, Facultad de Ciencias, Universidad de la República, Montevideo, Uruguay

**Keywords:** segmentation, anteroposterior patterning, regeneration, tapeworm, Wnt, Hedgehog

## Abstract

Tapeworms are parasitic flatworms that lack key features conventionally used to define the head-tail axis in free-living organisms, resulting in long standing questions regarding the true orientation of their main body axis. As adults, most tapeworms also exhibit a segmented body which has been considered an adaptation unique to the group. Anteroposterior (AP) patterning in free-living flatworms is controlled by β-catenin-dependent Wnt signalling and positional control genes are expressed by their musculature in highly regionalised domains. Here we investigate the expression of Wnt and Hedgehog components during the strobilar phase of the tapeworm life cycle during which larval tissues are lost and replaced through the continuous production of new tissues. Results reveal previously unidentified centres of signalling associated with their neuromuscular system and show that segments are marked by secondary, AP axes in agreement with the polarity of the primary body axis. Neuromuscular expression and communication between Wnt and Hedgehog signalling is consistent with embryonic and regenerative growth in planarians and is a common mechanism for establishing AP-polarised boundaries in a diverse range of segmented animals. Taken together our results suggest that segmentation in tapeworms represents a modified form of posterior regeneration, a common feature among flatworms.

## Background

Tapeworms are a medically and economically important group of helminth pathogens and one of the oldest recognised forms of parasitic worm [1]. Their segmented, or strobilar, adult body plan is a derived feature not only among flatworms (phylum Platyhelminthes) but also among members of the class Cestoda in which the repetition of body parts appears to have evolved in a step-wise fashion under selection for increased fecundity [2]. Their multi-host life cycles and highly derived form lacking most cephalised structures as well as a gut, has made their body plan difficult to homologise with other animals [3–5], leaving longstanding questions regarding the true polarity of their anteroposterior (AP) axis [6–8], the individuality of their segments [9] and more generally whether the mechanisms that underlie segmentation resemble those of other animals [10,11].

Anteroposterior (AP) patterning in free-living planarian flatworms is regulated by canonical, β-catenin-dependent Wnt signalling [12] and expression of ligands, inhibitors and receptors show distinct regionalisation along the main body axis [13]. These and other region-specific markers have been referred to as positional control genes (PCG) and are expressed continuously by muscle cells throughout their lives [14,15], maintaining an axial cartesian coordinate system [16] that is instructive to the somatic stem cells (neoblasts) responsible for all cellular renewal during growth, homeostasis and regeneration. Canonical Wnt signalling is a universal form of cell-cell communication in Metazoa with a conserved function in AP patterning through the combination of posterior Wnt signalling and anterior Wnt inhibition [17].

Extending this model to parasitic flatworms, Koziol et al. [5] investigated the expression domains of homologs of planarian Wnt components [18] and other canonical markers of AP patterning during larval metamorphosis in two tapeworm species with highly divergent morphologies. For the first time this provided a common basis of comparison and showed that despite gross morphological differences in the structure of their larvae, in both forms the site(s) of scolex formation (the ‘head’ containing the holdfast structures and CNS) is preceded by expression of anteriorising Wnt inhibitors, whereas the larval cyst tissues, which have evolved into a diverse range of morphologies in different tapeworm groups [19], express canonical, posteriorising Wnts. This resolved the question of the true developmental AP polarity of the larval worm and provided support for the idea that metamorphosis represents the phylotypic stage in their ontogeny with AP-regionalised patterns of Wnt expression mirrored across larval forms of tapeworms and in planarians [5]. It also demonstrated PCG expression in muscle cells and provided an explanation for their maintenance as a source of PCG expression in the otherwise non-motile cyst tissues. However, it left open the role of Wnt signalling during adult development, including questions regarding the polarity of the individual segments [7,8].

Metamorphosis of the tapeworm oncosphere results in the genesis of an encysted, juvenile worm with a fully mature scolex and rudimentary body that includes muscle, nerve and osmoregulatory systems (Fig. 1). In most tapeworms, sexual and strobilar development are then repressed until the larva is transmitted trophically to the enteric system of the vertebrate, final host. Following ingestion by the host, larvae excyst in the stomach and slough off their cyst tissues before becoming established in the small intestine. The excysted juvenile worm thus consists of little more than the scolex and adult development commences via elongation of the body, with the germinative neck region and strobila intercalated between the scolex and the opposing end of the worm (Fig. 1A). Sexual development is coupled to the segmental process and results in hermaphroditic sets of reproductive organs (proglottids) that are semi-compartmentalised through partial segmentation of the internal, medullary region of the worm. Adult development thus represents the second major transformation in their post-embryonic ontogeny and a significant departure from the typical, non-segmented flatworm body plan.

**Figure 1.**
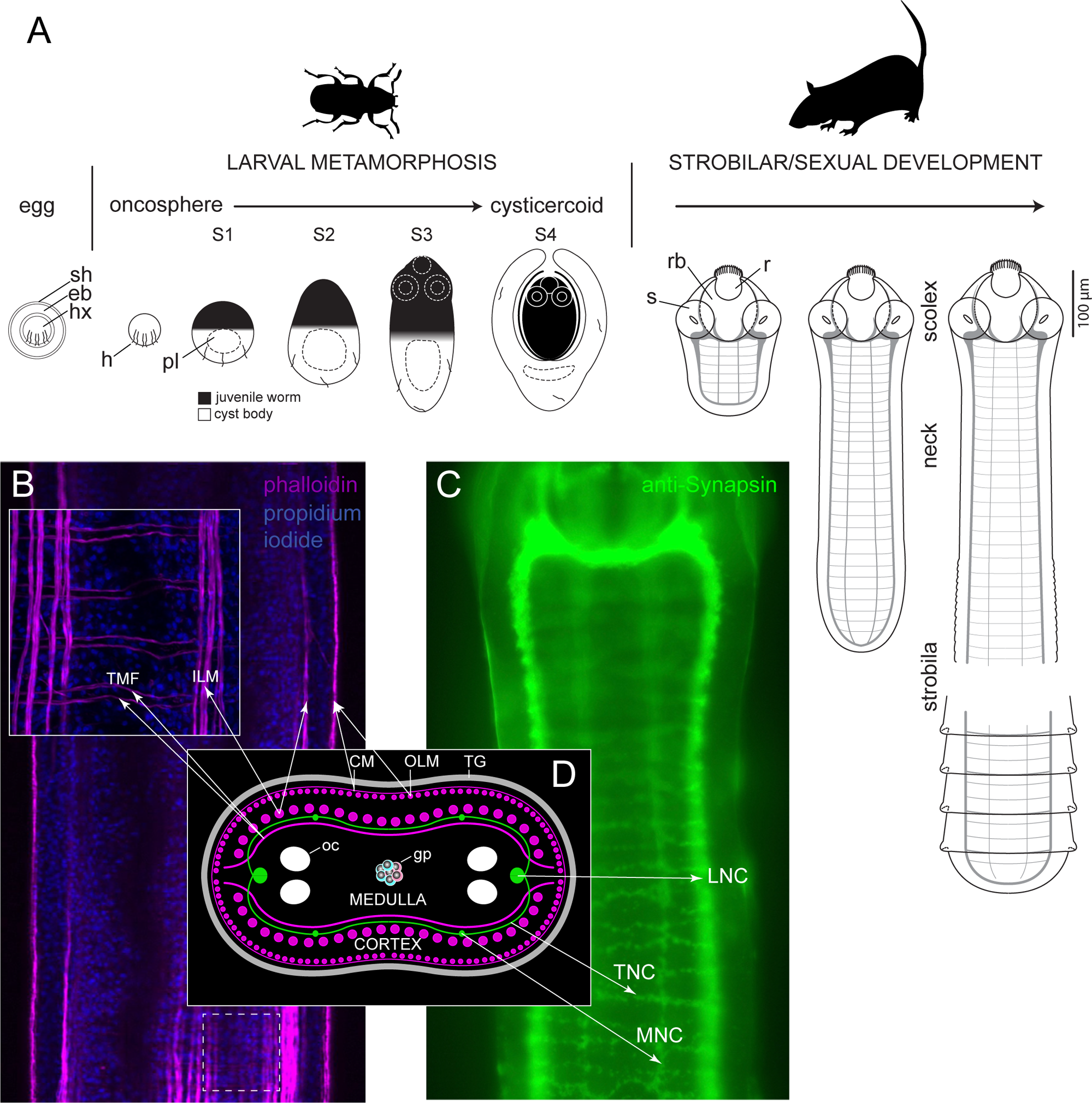
Ontogeny and neuromuscular anatomy of the mouse bile-duct tapeworm *Hymenolepis microstoma*. **A.** Larval metamorphosis and strobilar growth depicting the major stages of larval development and the three traditionally recognised regions of the strobilar adult (n.b. illustrations drawn to scale save the strobila detail which is significantly reduced in size relative to the neck/scolex). The life cycle is perpetuated when eggs expelled into the environment with the faeces of mice are consumed by grain beetles (e.g. *Tenebrio* spp.), releasing the oncospheral larvae that use their hooks to penetrate the intestine and enter the haemocoel where they take a week to metamorphosis into encysted, juvenile worms called cysticercoids [5]. Strobilar/sexual development is delayed until infected beetles are ingested by mice, releasing larvae that excyst in the stomach and slough their cyst tissues before entering the small intestine. There the juvenile worms, which consists of a mature scolex and rudimentary body, undergo elongation to form a neck region that generates the genital primordia and nascent strobila before the worms locate permanently in the mouse bile-duct. After two weeks the worms reach their maximum size and possess ∼650 segments (see [20]). Major elements of the neuromuscular system shown by phalloidin (**B**) and anti-Synapsin (**C**) immunostaining. Confocal imaging of the neck region of phalloidin stained worms shows the major muscle layers and development of paired, transverse muscle fibres (TMF) that define intersegmental boundaries (inset shows boxed region). Fluorescent microscopy of anti-Synapsin staining shows the primary elements of the nervous system consisting of the main longitudinal (LNC), medial longitudinal (MNC), and transverse (TNC) nerve cords. **D.** Diagram showing the neuromuscular anatomy of the neck in cross section. Abbreviations: eb, embryophore; CM, circular muscle layer; gp, genital primordia; h, hook; hx, hexacanth (= oncosphere) larva; ILM, inner, thick longitudinal muscle layer; oc, osmoregulatory canal; OLM, outer, thin longitudinal muscle layer; pl, primary lacuna (= cavity); r, hooked rostellum; rb, rostellar bulb; s, sucker; sh, membranous shell; TG, syncytial tegument; TMF, segmentally distributed, medullary transverse muscle fibres.

Here we extend investigations of PCG expression in tapeworms to the strobilar phase of the life cycle of the mouse bile-duct tapeworm *Hymenolepis microstoma* [20]. This classical mouse/beetle-hosted laboratory model has been the subject of a series of genomic [21–24], transcriptomic [25,26] and developmental studies [5,27] that underpin its utility as a contemporary model and is now supported by the first fully complete, chromosome-level genome assembly of a spiralian [28]. We use single and double fluorescent whole-mount in situ hybridisation (FISH) to examine the expression domains of genes encoding Wnt ligands (*wnt1*, *wnt11a*, *wnt11b*), inhibitors (*sfrp, sfl, notum*) and receptors (*fz4*, *fz5/8*, *fz1/2/3/6/7*) involved in the canonical Wnt pathway, as well as of a gene encoding a Hedgehog ligand, an upstream regulator of Wnt signalling in planarians [29]. Results show that the scolex and neck region are characterised by Wnt inhibition, while the transition to strobilar growth is marked by polarised expression of Wnt ligands and inhibitors in agreement with the polarity of the primary head-tail axis. We identify a Wnt11 paralog whose expression is spatially and temporally restricted to the initial formation of segments, suggesting it may have acquired a strobilation-specific role. Expression patterns reveal signalling centres associated with the neuromuscular system that link neural expression of *hedgehog* with muscular expression of *wnt*. Finally, we show that in adult tapeworms some of the muscle cell nuclei (myocytons) are uniquely clustered along the dorsoventral (DV) midline adjacent to the main longitudinal nerve cords (LNC) and are major foci of PCG expression. Taken together, our results suggest that the strobilar tapeworm body plan represents a modified form of flatworm posterior regeneration involving repeated specification of secondary AP axes.

## Results

### Spatial distribution of myocytons indicates their participation in segmentation and PCG expression

Given the known expression of PCGs by muscle cells in planarians [14] and larval tapeworms [5], we investigated the spatial distribution of muscle cells in adult *H. microstoma*. Unlike other segmented organisms, most of the musculature of tapeworms is not segmentally organized [4]. Their musculature includes a subtegumental muscle layer comprised of thin, outer circular and longitudinal muscle fibres, an inner layer of thick longitudinal bundles of fibres that attach to the apical rostellum (that by convention define the cortical/medullary boundary; Fig. 1D), and numerous dorsoventral fibres that run through the medullary region of the worm. At segmental boundaries there are AP-paired strands of transverse muscle fibres (Fig. 1B) that are dorsoventrally symmetrical and attach to the outer longitudinal muscle layer and traverse the medullary region just inside of the inner longitudinal muscle layer (Fig. 1D). These transverse fibres first develop in the anterior of the neck and later come to define the segmental boundaries. To the best of our knowledge, these are the only muscle fibres with a segmentary distribution along the AP axis.

The nuclei of flatworm muscle cells are located in non-contractile cell bodies called myocytons that are offset from the contractile fibres to which they are connected via thin cytoplasmic processes (see Fig. 3F in [14]). We analysed the spatial distribution of myocytons in adult worms by WMISH using probes for two Tropomyosin isoforms (*hm-tpm1*, HmN_000188900.3; *hm-tpm2*, HmN_000471300.3) [30,31] that produced identical results, as well as an isoform of Collagen (*hm-collagen*, HmN_000398500) [14]. We identified several different myocyte populations according to their spatial distribution (Fig 2). In the cortex we identified myocytons associated with the outer muscle fibres immediately beneath the tegument (Fig. 2A); myocytons of the inner longitudinal muscle layer (Fig. 2A’); and uniquely clustered myocytons at the dorsoventral midplane, external to the main LNCs that form ribbons along the margins of the worm and are hereafter referred to as ‘marginal myocytons’ (arrows in Figs. 2A’’ and B). The clusters are also paired by a smaller domain of expression on the opposite (medullary) side of the LNCs, seen as internal ribbons along the AP axis (arrowheads in Fig. 2B). In the centre of the medullary region, between the osmoregulatory canals, is another prominent domain of expression (arrowhead in Fig. 2B) representing muscle cells that presumably intercalate between the cells that give rise to the genital primordia, the precursors to the hermaphroditic proglottids [32,33] (Fig. 2B). For some of the myocyton populations in the inner muscle layer we could detect *tropomyosin* signal not only in the myocytons but also at the periphery of the muscle fibres, evidenced by double detection with antibodies against muscle tropomyosin isoforms [30]. However, in the marginal and cortical myocytons, it was not possible to unambiguously identify their associated contractile fibres.

**Figure 2.**
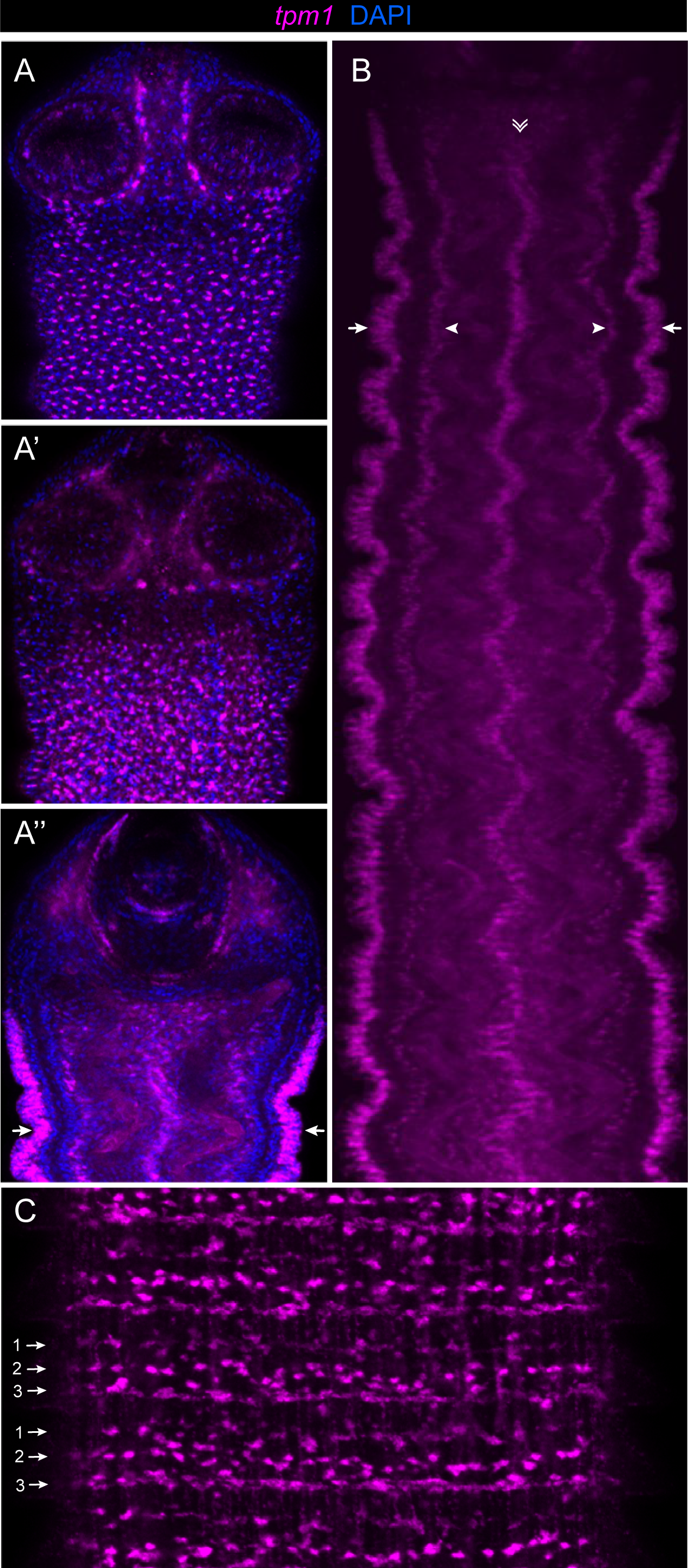
*Tropomyosin* expression reveals the spatial pattern of muscle cell nuclei. *Hm-tpm1* expression shows the punctate pattern of myocytons throughout the muscular system. Separate populations can be seen in association with the outer (**A**) and inner (**A’**) muscle layers, while a section through the dorsoventral midline (**A’’**) highlights the clustered arrangement of the myocytons (arrows) exterior to the main longitudinal nerve cords. Midline section through the neck region (**B**) shows the marginal myocytons (arrows) and associated internal ribbons of myocytons (arrowheads) that flank the main LNC, and a central ribbon of foci (double arrowhead) that represent myocytons presumably intercalated between cells that give rise to the genital primordia. In maturing segments (**C**) some myocyton populations come to be arrayed in three transverse rows (arrows) that mark the positions of the three transverse nerve rings that develop in each segment (see Fig. 1A).

**Figure 3.**
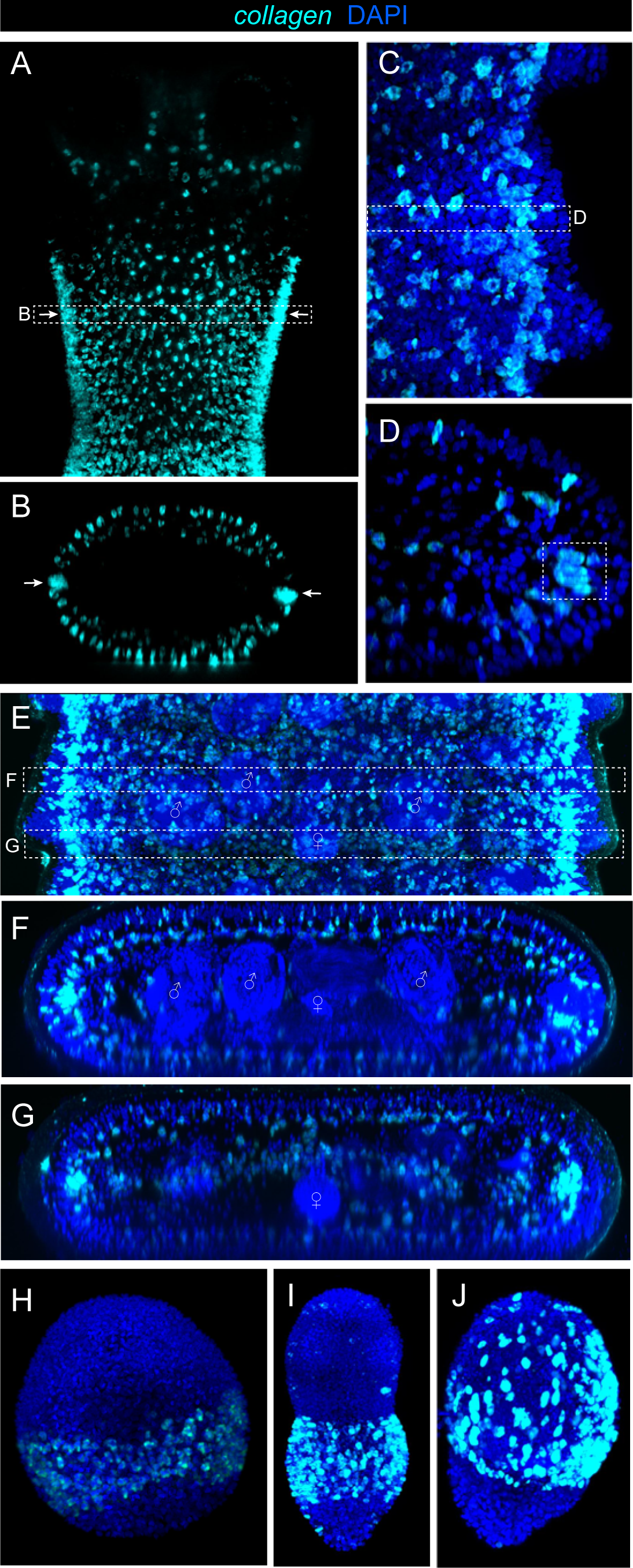
*Collagen* shows PCG-like expression in muscle cells. *Hm-collagen* is expressed by a subset of the myocyton populations seen via expression of *hm_tpm1* (Fig. 2) and is restricted primarily to the cortical region. In the scolex (**A**) there are few *collagen*+ cells whereas the punctate pattern of the muscle layers, together with clustered pattern of myocytons at the margins (arrows in **A-B**) begins in the neck. **C-D**. Higher magnification of segment margins shows the clustered myocytons at the margins (box in **D**). **E-G**. Expression in mature segments. Transverse reconstruction through the level of the testes (**F**) shows punctate expression corresponding to the two muscles layers of the body wall, whereas reconstruction at the segmental boundary (**G**) shows additional foci running through the centre of the worm. **H-J**. Expression during larval development (see also Additional files 2-3). **H**. Stage 1 larva showing a band of expression in the posterior hemisphere, corresponding to the region of the cyst cavity (primary lacuna; see Fig. 1). **I**. Stage 3 larva showing marked differentiation of the anterior (nascent worm) and posterior (cyst) regions. **J**. Stage 4 encysted larva showing expression in the cyst tissues that surround the juvenile worm.

Unexpectedly, we found that although the inner longitudinal muscles run continuously through the strobila, some of their myocytons become intra-segmentally distributed along the AP axis as segments mature, resulting in the presence of three transverse ‘stripes’ of myocytons (Fig. 2C) that correspond spatially to the three nerve rings present in mature segments (Fig. 1; [4,34]). The distribution of these myocytons, as well as those of the transverse muscle filaments that define segmental boundaries, suggests that muscle cells could participate directly in the segmentation process, or that their precursors could be among the earliest cells to respond to signals regulating segmentation.

We also investigated the spatial patterns of myocytons via expression of *collagen*, as muscle cells have been shown to be the main cell type expressing diverse extracellular matrix components in the parenchyma of planarians [35]. *Hm-collagen* was found to be expressed by a subset of the myocyton populations expressing *hm-tpm1/2* and was associated primarily with the main muscle layers (Fig. 3). Expression of *collagen* in the medullary region, which contains numerous dorsoventral fibres, was weak except at segmental boundaries where a line of internal foci was seen along the DV midline (Fig. 3G). Little expression was seen in the scolex (Fig.3A), whereas the outer and inner muscle layers and the ribbons formed by the marginal myocytons showed strong expression. During larval metamorphosis *collagen* expression was restricted to the posterior hemisphere in the region of the primary lacuna (Figs. 3H-J, Additional files 2-3). Although its expression underrepresents the total myocyton population, its more restricted pattern is instructive in including many of the prominent domains of the Wnt components described below.

### Wnt inhibitors are expressed in the scolex and neck

We found that the scolex and neck are characterised by the expression of secreted frizzled receptor (sFRP) genes (Fig. 4) which are inhibitors of canonical Wnt signalling [17]. Tapeworms have two sFRP genes: a homolog of sFRPs that canonically act to specify the anterior pole in metazoans (*hm-sfrp*; HmN_000556500) and a *sfrp*-like paralog (*hm-sfl*; HmN_000359400), both of which were shown previously to be expressed in anterior domains in larval worms [5]. In adult worms, *sfrp* is expressed in a punctate pattern in the outer, cortical tissues of the neck, before being abruptly down-regulated at the start of the strobila where external segmentation first becomes visible (Figs. 4A-C). A number of internal foci are also seen in the scolex and confocal reconstruction of *sfrp* together with anti-Synapsin staining [34] shows that these are associated spatially with the main cerebral ganglia and its terminal branches that innervate the holdfast structures (Figs. 4D-E). In the neck region *sfrp* is expressed only in the outer cortex, corresponding to the outer longitudinal and/or circular muscle layers, and in the marginal myocytons described above (Figs. 4A-F).

**Figure 4.**
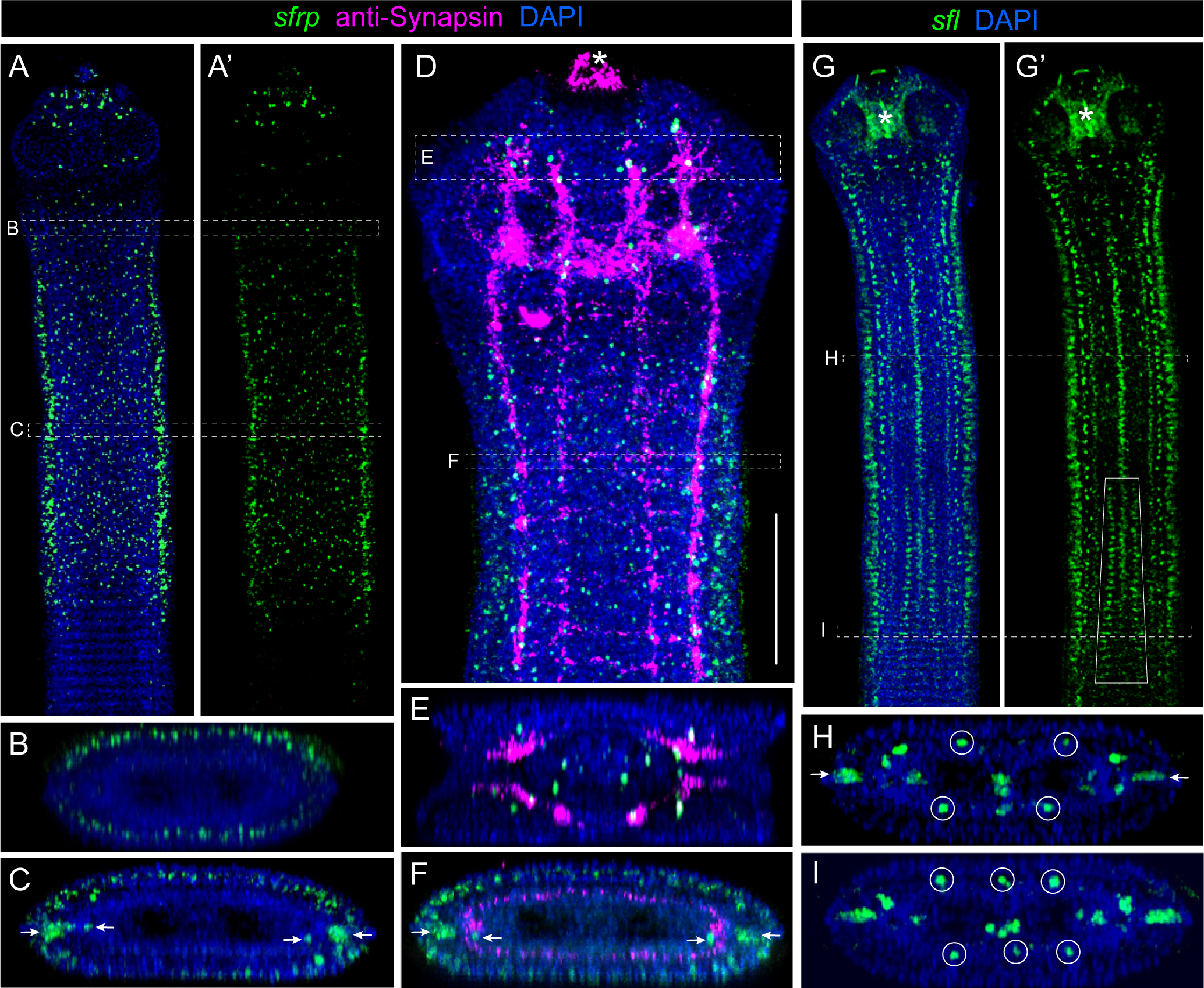
Expression of *sfrp* and *sfl* mark the scolex and neck as wnt inhibitory. **A.** *Hm-sfrp* expression in the scolex and neck shown with (**A**) and without (**A’**) DAPI nuclear staining. **B.** Transverse reconstruction through the neck showing expression restricted to the outer cortical tissues, and further down (**C**), additional large foci of expression at the DV midline immediately exterior to the main LNCs (the ‘marginal myocytons’; see text), with smaller foci on opposing sides of the LNCs (arrows). Combined with anti-Synapsin staining (**D-F**; see also Fig 1) shows expression in the scolex is restricted to internal foci associated with the major elements of the nervous system (**E**), whereas reconstruction through the neck region (**F**) shows the position of the peripheral nervous system at the cortical-medullary boundary (see also Fig. 1D). **G-I.** In contrast, expression of *hm-sfl* in the neck (**G**-**G’**) shows internal (medullary) expression. Transverse reconstruction through the neck (**H**) shows internal foci associated with the nervous system, including the marginal myocytons (arrows) and junctions of the medial longitudinal and transverse nerve cords (circled), as well as expression in the central area where the genital primordia develop. The boxed area in G’ shows the ‘signalling quartet’ (see text) that appears in the transition from the neck to strobila and reconstruction through the immature strobila (**I**) shows these foci together with the new central foci of expression (circled). Asterisks in panels **D**, **G** and **G’** indicate artefactual signal in the glandular, rostellar bulb (see also [34]).

*Hm-sfl* is also expressed in the scolex and neck regions but differs from *sfrp* expression in most other respects. In particular, it is also expressed segmentally and thus in all regions of the worm save the primary posterior pole represented by the terminal tissues (Figs. 4G-I). It also differs in being expressed in the medullary region rather than the cortex, with the exception of the marginal myocytons in which both genes are expressed. Medullary expression includes central and lateral ribbons of cells along the DV midline that mirror the pattern of *hm-tpm* (Figs. 2A”, B). In the neck region there is also a repeated pattern of internal foci (Fig. 4H; see also Fig. 10D’) that correspond to the junctions of the dorsal and ventral medial and transverse nerve cords (Fig. 1C). Unlike the muscular system which exhibits a segmental pattern of transverse, medullary fibres starting in the neck, the peripheral nervous system (PNS) in the neck and into the early strobila is not segmental, and in contrast exhibits a basic orthogonal pattern typical of flatworms [36] before the pattern eventually comes into register with the segmental boundaries of maturing segments. Unlike the other domains of *sfl* that mirror the distribution of myocytons, these domains are consistent with expression by the nervous system and we later show that *sfl* and *hedgehog* co-localise at these nerve junctions (Fig. 10D-E).

In the transition from the neck to the strobila a new, internal quartet of foci appears, hereafter referred to as the ‘signalling quartet’ (Fig. 4I; Additional file 4). This quartet of *sfl* foci arises in register with the early segmentation of the strobila, with two dorsal and two ventral cells in each nascent segment. As the strobila grows, the position of the signalling quartet is seen to widen toward the junctions of the medial and transverse nerve cords (see boxed area in Fig. 4G’ and Fig. 5B). Furthermore, an additional segmentary domain of expression appears in the dorsal and ventral medial lines. Hence the transition from neck to strobila is characterised in part by new domains of *sfl* expression along the AP axis.

**Figure 5.**
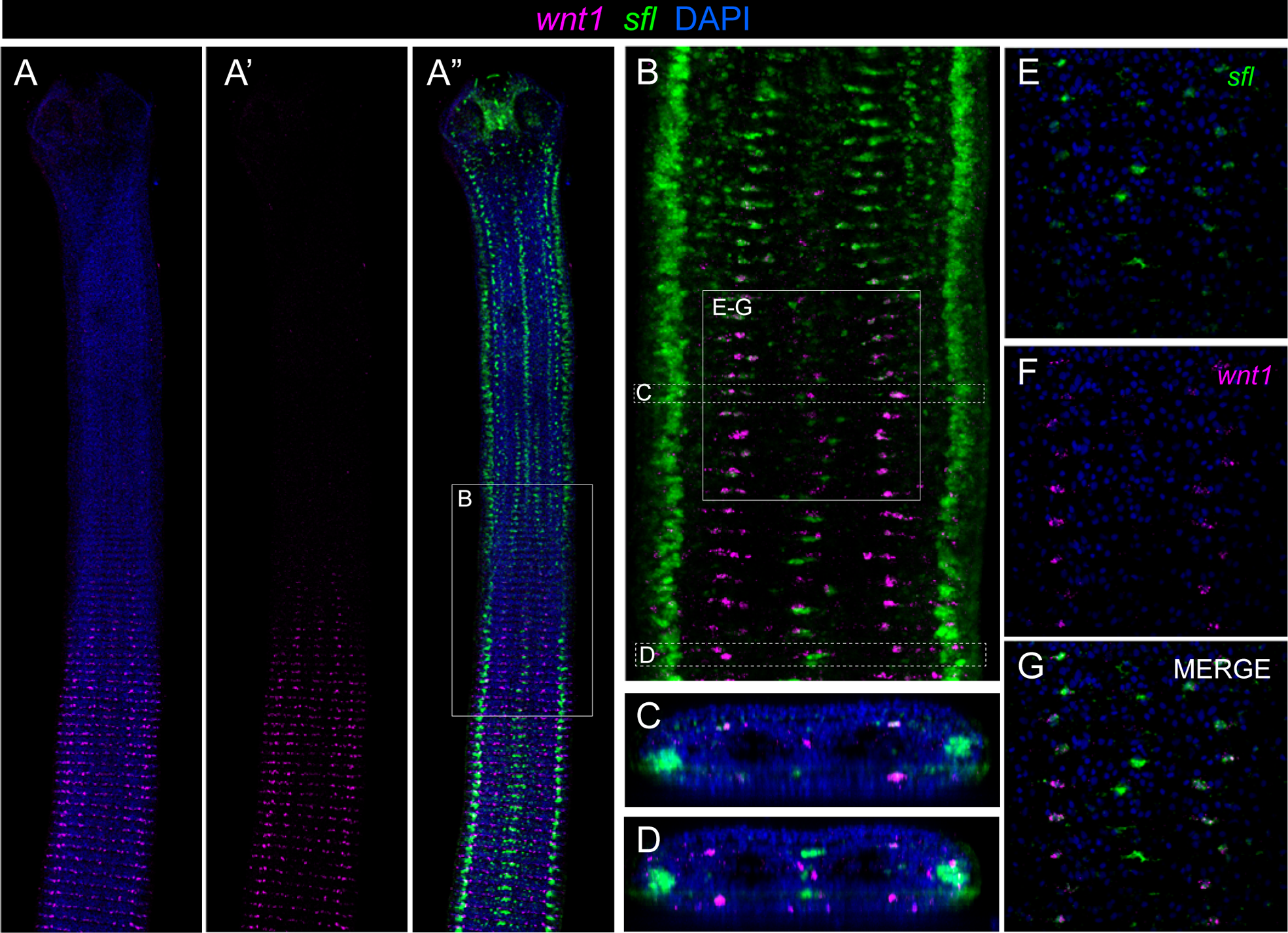
Initial expression of the canonical posterior *wnt1* gene shows co-localisation with the inhibitor *sfl*. **A-A”.** *Hm-wnt1* expression shown with (**A**) and without (**A’**) nuclear staining, and in combination with the inhibitor *sfl* (**A”**). Wnt1 expression begins in the area where the neck transitions to the strobila (boxed in **A”**) which is imaged at higher magnification in (**B**). From the anterior (top), expression is seen first in the genital primordia, then in the signalling quartet (see text) and subsequently in stripes along the leading edge of the nascent segments. In the signalling quartet of cells *wnt1* and *sfl* expression is initially seen to co-localise (**C, E-G**), while separate foci appear later as segments form. In the genital primordia, *wnt1* and *sfl* are expressed by separate populations of cells (**B, D**).

### Strobilar development involves AP-polarised expression of Wnt inhibitors and ligands

The first instance of posterior Wnt gene expression during strobilar growth begins at the transition between the neck and strobila (Fig. 5). *Hm-wnt1* (HmN_000328000), a homolog of the Wingless/Int gene family, becomes visible as a gradient in the signalling quartet and associated foci described above (Fig. 5). Double FISH shows that *wnt1* initially co-localises with *sfl* in the signalling quartet (Fig. 5E-G’) but not in the central domain in which separate foci are evident (Fig. 5D). As *wnt1* expression increases, *sfl* diminishes in the signalling quartet and is replaced by *wnt1*, while in the central foci *sfl* expression increases (Figs. 5A”, B, D; Additional file 5). In the strobila *sfl* expression is again seen in a quartet pattern (Fig. 6B) and in nascent segments the domains of both *sfl* and *wnt1* increase around the circumference of the worm, eventually forming AP-polarised stripes at segmental boundaries (Fig. 6; Additional file 6). Opposing expression of *sfl* and *wnt1* thus marks each segment with its own secondary AP axis in agreement with the polarity of the primary body axis. As segments mature, *sfl* expression also becomes visible along the DV midline at segmental boundaries (Figs. 6D, G) and its expression in the marginal myocytons forms a gradient that diminishes posteriorly (Fig. 6F, F’). In contrast, *wnt1* expression is notably absent from both the marginal myocytons and medullary musculature (Figs. 6E, H).

**Figure 6.**
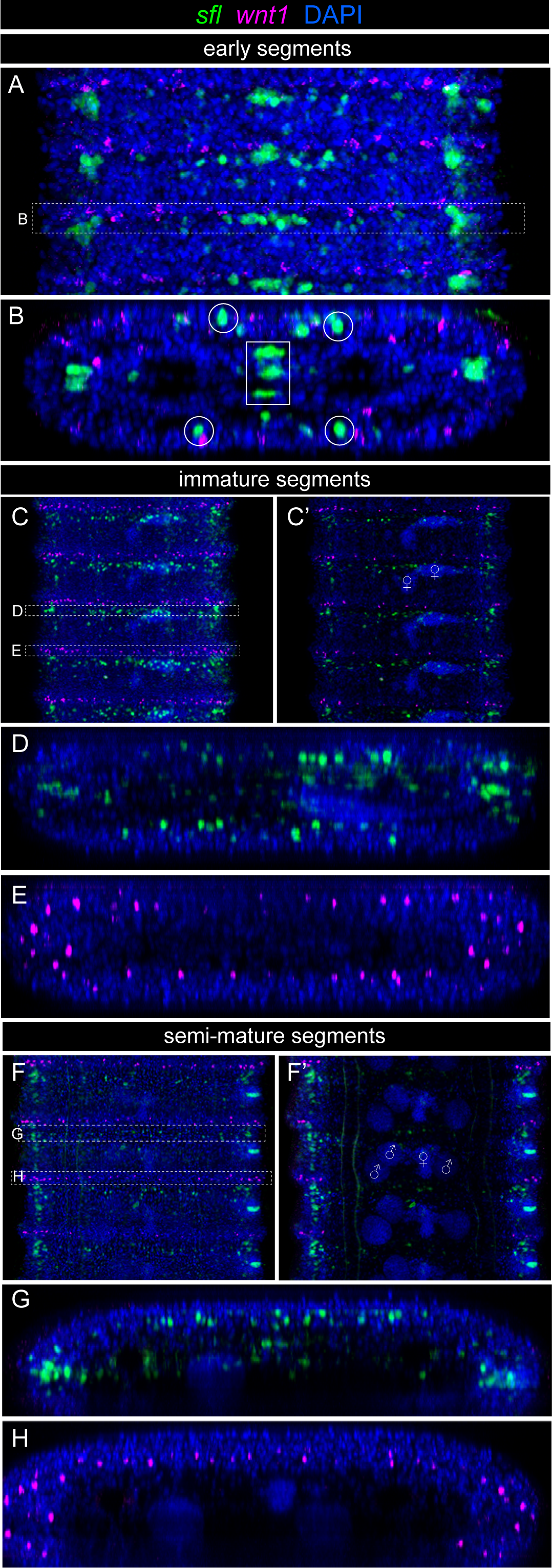
*Wnt1* and *sfl* show AP-polarised expression at segmental boundaries. **A.** Early segments show AP-polarised expression of *hm-sfl* and *hm-wnt1* at segmental boundaries. **B.** Confocal reconstruction shows the quartet of *sfl*-expressing cells (circled) and three dorsoventrally-stacked cluster of cells (box). Immature and semi-mature segments are shown as maximum projections of the entire image stacks (**C, F**) and of only the central portion of the worms (**C’, F’**). **C.** Immature segments show *sfl* expression marking the anterior (toward scolex) and *wnt1* the posterior boundaries of each segment. **C’.** Internal sections make visible the seminal receptacle and central ovaries (11) and nascent testes. **D-E.** Confocal reconstructions through the segmental boundaries (as indicated in panel **C**) shows that expression *sfl* (**D**) and *wnt1* (**E**) expands to encircle the segments, and that *sfl* is also expressed in cells that run through the middle of the worm at the segmental boundaries (**D, G**), whereas *wnt1* expression is restricted to the cortical tissues and is not expressed in the medullary region (**E, H**). In more mature segments (**F-H**), clusters of *sfl*-expressing cells can be seen arrayed along margins the of the worm at the DV midline (**F’**) forming a gradient diminishing posteriorly before being interrupted by a narrow band of *wnt1* expression. Internally, testes (11) and ovaries (11) are visible via DAPI staining, while strong expression of *sfl* is also seen in the area of the genital pores which open at the dextral margins of the worm. Confocal reconstructions (**G-H**) through the areas indicated in panel **F** show the patterns described above, with *sfl* expressed not only in cortical tissues, but also internally through the central axis of the worm, and in the large marginal clusters of myocytons (see text), whereas *wnt1* expression is cortical and restricted to a narrow transverse plane, and is not expressed in the marginal myocytons.

### Expression of *wnt11a* marks the transition to strobilar development

The transition between the neck and strobila forms a morphological gradient marked by crenulations of the outer body wall that form in register with the internal segmentation. Strikingly, this transition is tightly demarcated by the expression domain of one of two Wnt11 paralogs (*hm-wnt11a*; HmN_002147900) that starts prior to and includes the first visible signs of segment formation (Fig. 7, Additional file 7). *Wnt11a* begins as a gradient first in the central region and then as a series of segmental, punctate stripes, that appear in cross section as two concentric rings in the cortical tissues (Figs. 7D-E). Double FISH shows that its initial expression begins together with *wnt1* (Fig. 7C’) and then fades as *wnt1* expression increases and comes to be expressed as stripes. Expression of *wnt11a* is thus highly ephemeral and spans a region along the AP axis roughly equal in length to that of the neck (∼500 um). Results also show that it is expressed in cell populations distinct from those that are *wnt1+* (Figs. 7C-E). Double FISH with *sfl* shows that the onset of *wnt11a* expression is slightly after the initial change to segmental expression of *sfl* represented by the signalling quartet (Fig. 7F). Like *sfl*, *wnt11a* is also expressed in the marginal myocytons and in the central medullary region, but expression in the central region begins and ends earlier in development than the cortical expression, giving the appearance of an anteriorly offset ‘streak’ (Fig. 7F, Additional file 7).

**Figure 7.**
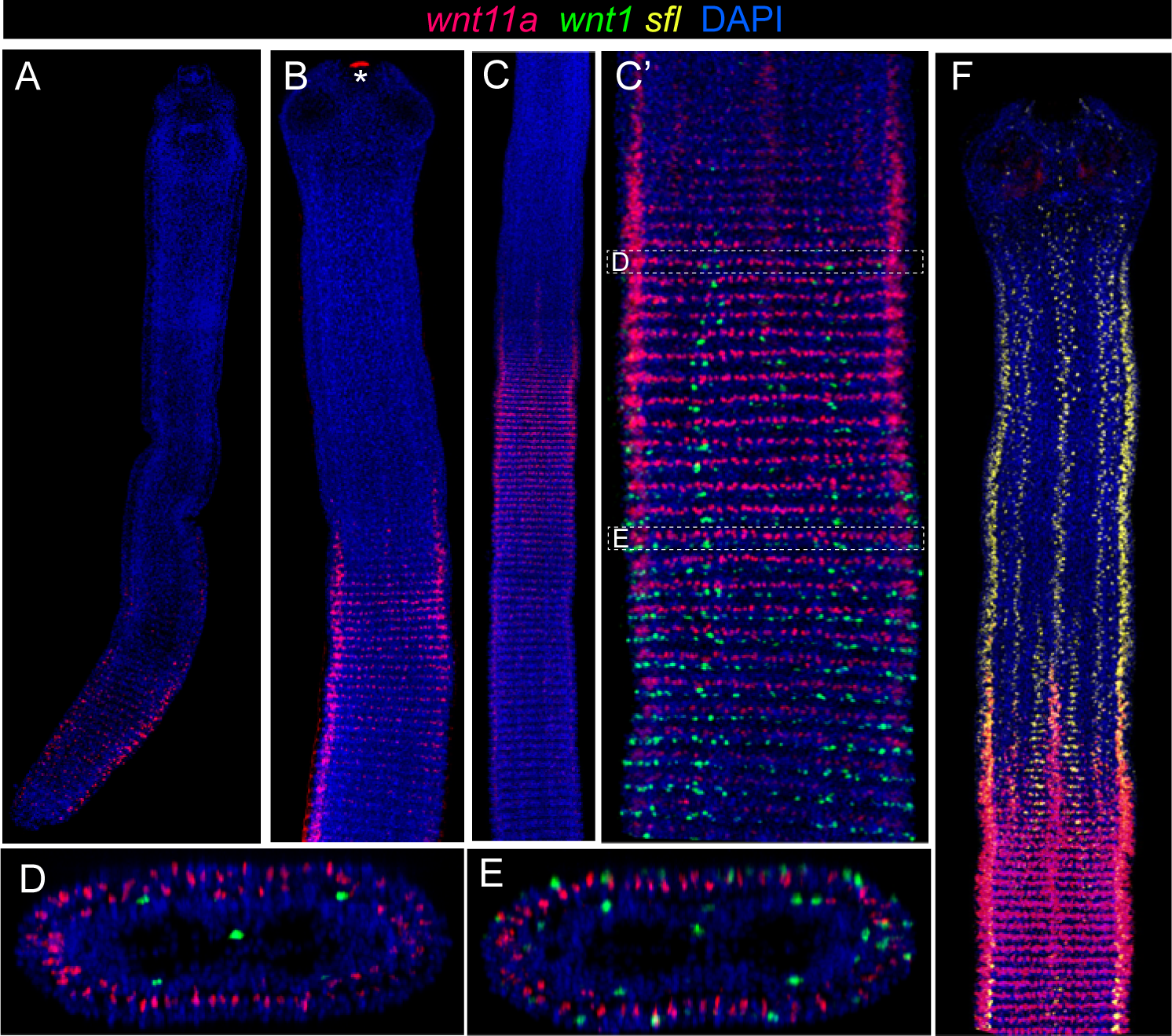
*Wnt11a* marks a transition zone between the neck and strobila. *Hm-wnt11a* forms a gradient of expression restricted to the ‘transition zone’ between the neck and strobila where segments become defined. *Wnt11a* expression in three-day-old, pre-strobilar juvenile worms (**A**) and in strobilar adults (**B-C**); see also Additional file 7. **C’.** Detail of *wnt11a* and *wnt1* expression in **C**. **D-E.** Transverse confocal reconstructions of **C’** showing cortical expression of *wnt11a* in two muscle layers, and early (**D**) and later (**E**) expression of *wnt1*. **F**. Expression of *wnt11a* combined with *hm-sfl* shows that *wnt11a* begins immediately after the pattern of *sfl* changes in the transition zone (see also Additional files 4-5). Asterisk in **B** indicates artefactual signal in the rostellum (see also Fig 4D).

### *Wnt11b* and *notum* expression is restricted to the strobila

Expression of the other Wnt11 paralog, *hm-wnt11b* (HmN_000022800), is similar in pattern to *wnt1* but starts abruptly in the strobila rather than forming a gradient beginning in the transition zone (Fig. 8). Like *wnt1*, *wnt11b* is restricted to the posterior segmental boundaries and becomes circumferential as segments mature, but differs in also being expressed by the marginal myocytons and across the DV midline at segmental boundaries (Figs. 8E, G), similar to *sfl* expression (cf. Fig. 6G) with which it is AP-polarised. *Wnt11b* also shows a few prominent foci that lie along left-right midline at segmental boundaries, bisecting the medullary region of the worm (Fig. 8C, E, G). Posterior segmental boundaries are thus marked by expression of the posterior Wnt genes *wnt1* and *wnt11b*, as well as by *wnt11a* in the transition zone.

**Figure 8.**
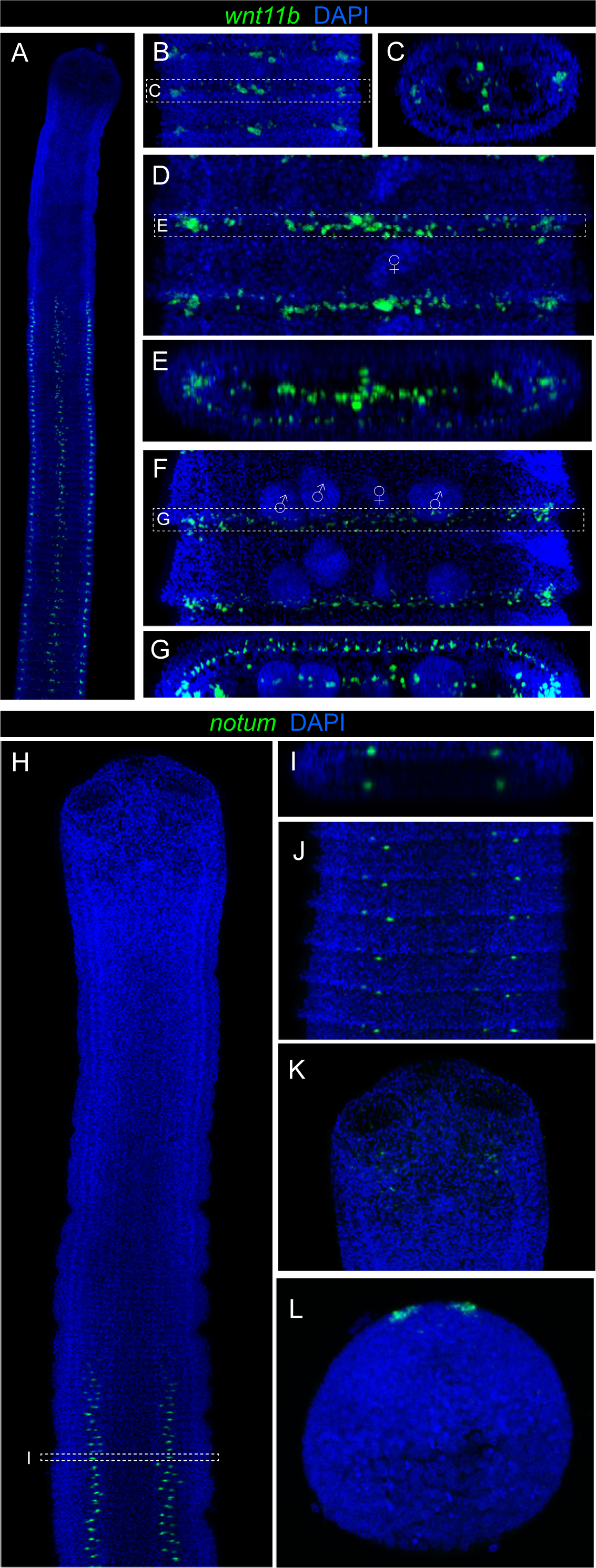
*Wnt11b* and *notum* expression is restricted to the strobila. **A.** *Hm-wnt11b* shows segmental expression restricted to a narrow, transverse plane at the posterior margins of the segments that begins abruptly post-transition zone. **B.** Early segments. **C.** Transverse reconstruction showing expression in the marginal clusters and three central foci (see also *Hm_wnt1*; Fig. 6B) that bisects the worm along the DV axis. Expression in immature (**D**) and mature (**F**) segments shows that foci increase in number to encircle the cortical region just outside of the inner longitudinal muscle layer. In addition, medullary foci expand across the dorsoventral midline forming a line of expression through the centre of the worm (disrupted by the osmoregulatory canals) (**E, G**). **H-L**. *Hm-notum* expression in adult and larval worms. **H**. *Hm-notum* expression is similar to *hedgehog* (see Fig. 10) in being expressed by the quartet (**I**) of cells at segmental boundaries (**J**), but differs in only being expressed after segments have been clearly defined. Expression in a single quartet of cells is also seen in the scolex (**K**). In Stage 1 larvae (**L**) *notum* is expressed in bilaterally symmetrical, apical clusters of cells at the anterior pole. Symbols: 11, seminal receptacle; 11, testes.

Like sFRPs, Notum is an inhibitor of canonical Wnt signalling [37] and *hm-notum* (HmN_000848900) expression (Figs. 8E-I) was found to be restricted to the nerve junctions (Fig. 8F) at segmental boundaries in the strobila, and immediately behind the suckers in the scolex (Fig. 8K). Like *wnt11b*, *notum* expression starts abruptly in the strobila after the initial formation of segments in the transition zone. As *notum* was not included in our previous study of Wnt signalling during tapeworm metamorphosis [5], we investigated its expression in larvae here and found that it is expressed anteriorly in two sub-apical, bilateral foci (Fig. 8I).

### Frizzled receptor gene expression is regionalised along the AP axis

Proteins of the Frizzled family function as receptors of Wnt ligands and like other components have distinct expression domains along the AP axis during embryogenesis [38]. We examined the expression patterns of three frizzled receptor genes with predicted roles in Wnt signalling (Fig. 9): *hm-fzd1/2/3/6/7* (HmN-000227100), *hm-fzd4* (HmN_000319700; [5]) and *hm-fzd5/8* (HmN-000386300). Homologs of *fz5/8* have a canonical role in specifying the anterior pole and are expressed in the anterior organizer of adult planarians [39]. We found weak expression of *hm-fzd5/8* as the base of the rostellum (Fig. 9A) but no expression in the neck region (Fig. 9B). In the strobila expression is first seen as a gradient in the marginal myocytons, the central region and in the signalling quartet (Figs. 9C, D), whereas in more mature segments anterior boundaries were marked by circumferential expression in the cortex and DV midline expression in the medulla (Fig. 9F), as well as a diminishing gradient of expression at the margins (Fig. 9E), mirroring the segmental expression pattern of *hm-sfl* (cf. Fig. 6).

**Figure 9.**
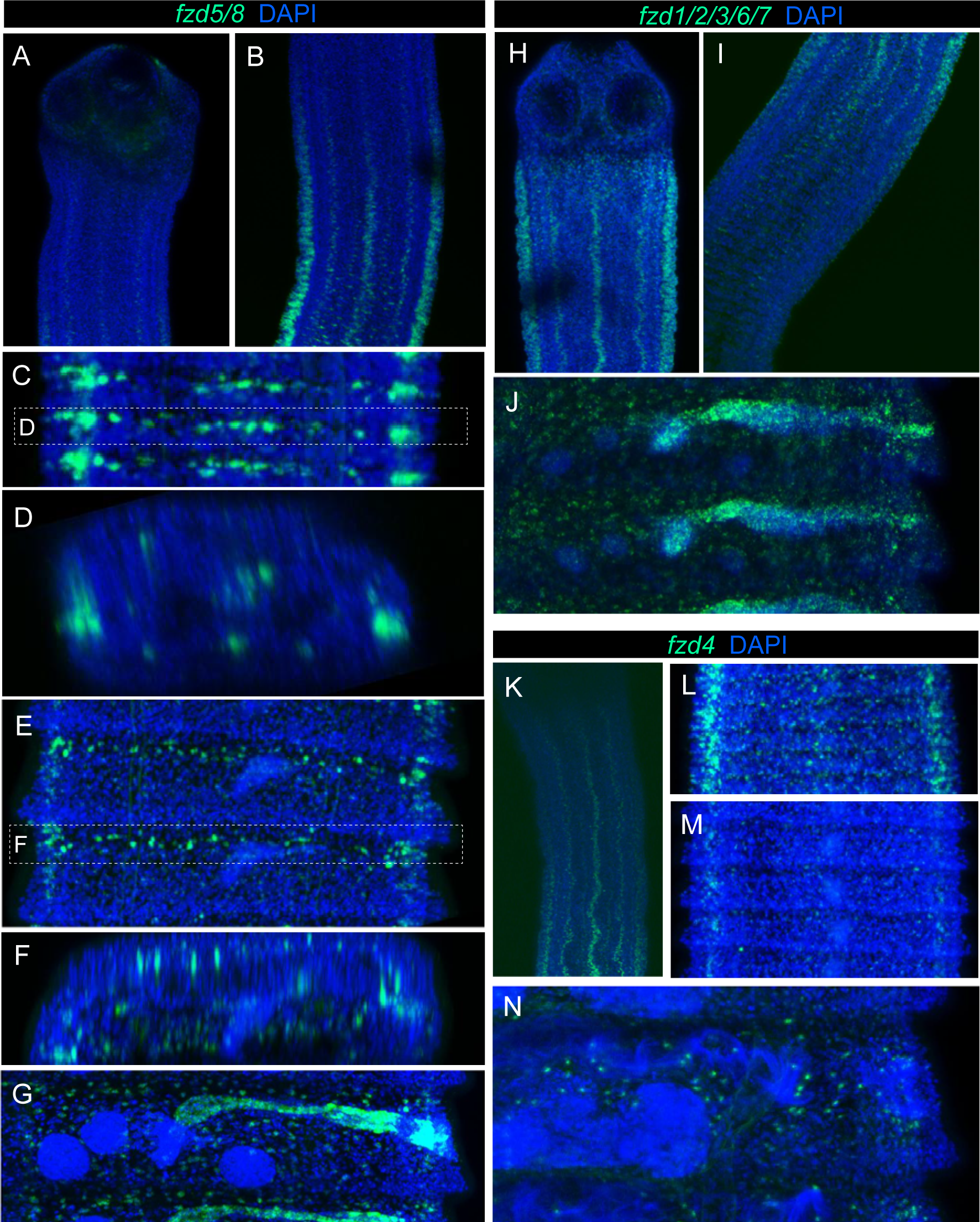
Frizzled receptor genes show regionalised expression along the AP axis. Frizzled receptor genes show similar domains of expression but differ in the regions of the worm they are expressed. **A-F.** *Hm-fzd5/8* shows weak expression in the scolex, but not in the anterior of the neck (**A**). Expression begins in a gradient nearer the transition zone (**B**) and continues throughout the strobila in a pattern similar to *hm-sfl* that includes the marginal myocytons, signalling quartet and central region (**C-D**). Maturing segments are marked by cortical expression at anterior margins (**E**) and by DV midline expression at segmental boundaries (**F**), also similar to *sfl* expression. In mature segments there is secondary, medullary expression seen as a punctate pattern throughout the parenchyma, as well as expression in the seminal receptacle, vagina, cirrus and genital pore (**G**). **H-J.** *Hm-fzd1/2/3/6/7* expression begins at the top of the neck (**H**) and then fades in the transition zone. The pattern of expression is similar to *fzd5/8*, including secondary, medullary foci in mature segments (**J**). **K-N**. *Hm-fzd4* is expressed in both the neck and strobila in similar domains to the other frizzled genes, but secondary expression associated with the reproductive ducts was not seen (**N**).

*Hm-fzd1/2/3/6/7* expression domains were similar to *hm-fzd5/8* but restricted regionally to the neck, beginning at the top of the neck where the marginal myocytons first appear (Fig. 9H). Its expression then fades in the transition to the strobila where *fzd5/8* expression begins (Fig. 9I). An identical pattern was described for the ortholog in *H. diminuta* (HDID_0000773501) that was identified in an RNA-Seq screen for genes enriched in the anterior of the neck region [40].

*Hm-fzd4* was expressed in both the neck and strobila and again showed similar domains to the other frizzled genes. However, unlike *fzd1/2/3/6/7*, expression in the marginal myocytons is absent from the top of the neck and begins as a gradient nearer the transition to the strobila (Fig. 9K). In the strobila its pattern was similar to *fzd5/8* in being expressed around the anterior of the segments (Fig 9L) and in a gradient at the margins (Fig. 9M). Frizzled receptor genes are thus highly regionalised along the AP axis and have expression domains that mirror the patterns of other components.

In maturing segments the frizzled genes also showed punctate expression in the medullary region (Figs. 9G, J, N) consistent with expression by myocytons of the dorsoventral muscle filaments. *Fzd5/8* and *fzd1/2/3/6/7* additionally showed expression in the seminal receptacle and vagina, the cirrus sac and the marginal genital pore (Figs. 9G, J).

### *Hedgehog* expression indicates a link with Wnt signalling via the nervous system

Hedgehog signalling has been shown to be an upstream promotor of Wnt signalling in flatworms [29] and we investigated *hedgehog* expression in larval and adult worms (Fig. 10). Combined with anti-Synapsin staining showed that in adults *hm-hedgehog* (HmN_000068600) is expressed in the scolex in discrete foci associated with neural ganglia (Fig. 10A). In the neck it is expressed in clusters of cells in the central, medullary region of the worm and at the junctions of the medial and transverse nerve cords (Fig. 10B) and both domains persist into the strobila (Fig. 10C). Double FISH shows co-localisation of *hedgehog* and *sfl* in both the nerve junctions (Fig. 10D) and central region (Fig. 10E). In the transition zone and early strobila expression in the nerve junctions remains consistent with the orthogonal pattern of the nervous system, which in turn remains independent of the underlying segmentary pattern of the medullary region (Fig. 10D”). Double FISH in larval worms showed that during metamorphosis *hedgehog* marks the sagittal midline on the dorsal and ventral surfaces, consistent with a canonical role in midline patterning [41], whereas *sfl* marks the dorsoventral midline at the margins (Figs. 10F-I; Additional files 8-11).

**Figure 10.**
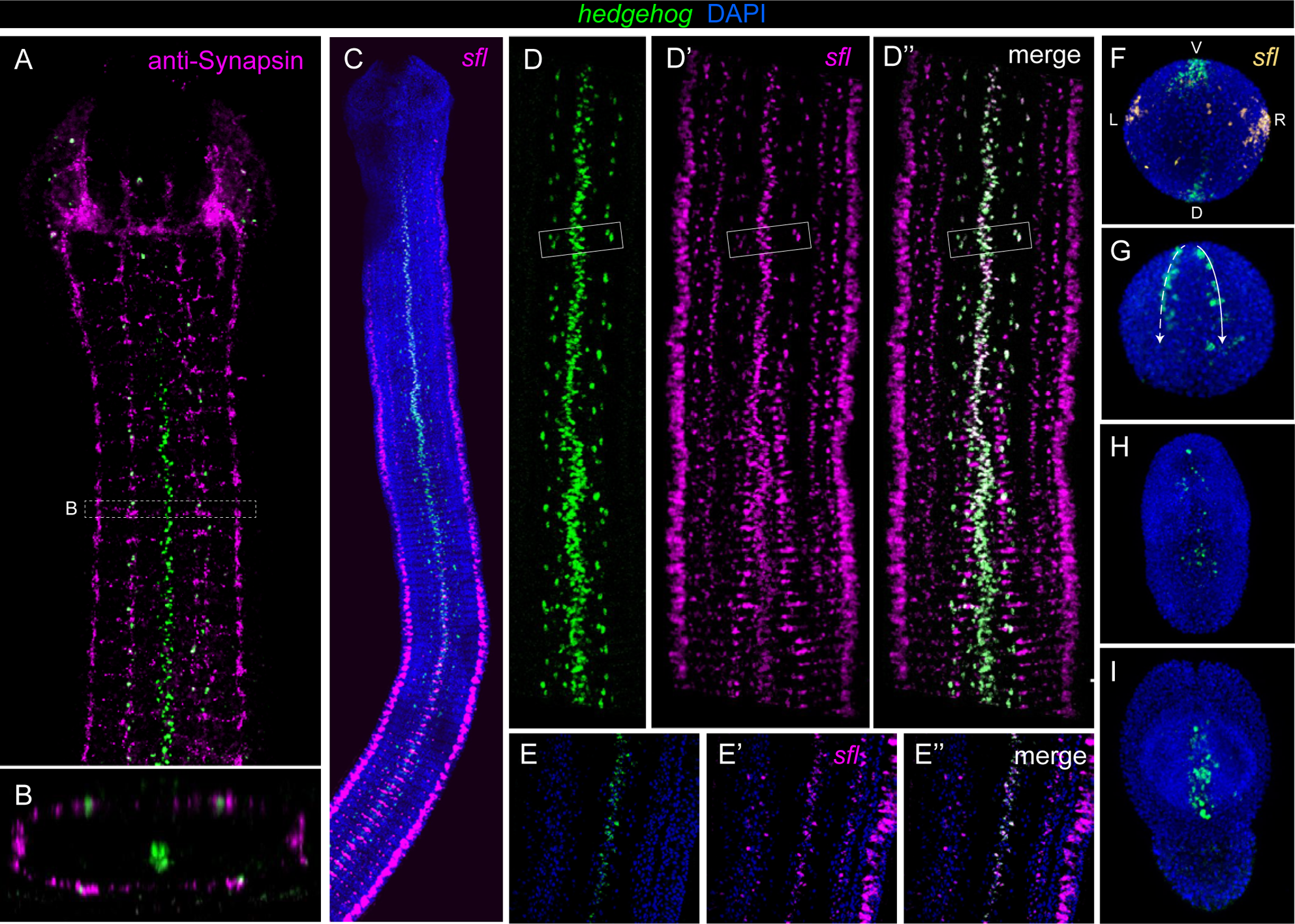
*Hedgehog* expression is associated with the nervous and reproductive systems. **A.** *Hm-hedgehog* expression in the scolex and neck combined with anti-Synapsin staining of the nervous system shows expression in the signalling quartet, corresponding to the junctions of the transverse and medial longitudinal nerve cords (**B**), as well as expression in the central, medullary region. Expression of *hedgehog* combined with *hm-sfl* (**C-F**) in the neck and transition zone shows co-localisation in the signalling quartet (**D-D”**, boxed regions show a single quartet pattern) and central region (**E-E”**). **F-I**. *Hedgehog* expression during larval development (see also Additional files 8-11). **F**. Apical view of a Stage 1 larva showing combined expression of *hedgehog* and *sfl* that demarcate the dorsal (D) – ventral (V) and left (L) – right (R) axes, respectively. **G**. Stage 1 larva showing lines of expression at the dorsal and ventral midlines of the anterior (i.e. worm forming) hemisphere. Continued expression in Stage 3 (**H**) and encysted, Stage 4 (**I**) larvae.

## Discussion

### Strobilar growth reveals centres of PCG expression associated with the neuromuscular system

The patterns revealed by *tropomyosin* and *collagen* expression encompass most of the expression domains of the Wnt ligands, antagonists and receptors, consistent with Wnt signalling being established and maintained by the musculature [14], which is most likely an ancestral system in bilaterians for encoding positional information via the body wall [42]. Unexpectedly, their expression revealed the clustered arrangement of the marginal myocytons which run continuously through the neck and strobila and are thus not overtly segmental in their arrangement. However, PCG expression in these clusters becomes discontinuous, segmental and AP-polarised. They are also positioned along the DV boundary which is known to act as an organising centre [13] and are immediately exterior to the main LNCs, allowing for close, cell-cell communication between the systems. We suggest that these uniquely arranged myocytons act as signalling centres, or organisers, that regionalise the AP axis during adult growth. Notably, this clustered pattern of myocytons is not observed in the musculature of planarians [14] and a broader survey of flatworms is needed to determine the extent to which the arrangement may be unique to *H. microstoma*, to tapeworms, or other flatworms.

The transverse muscle fibres that begin at the top of the neck (Fig. 1B) are the first overtly segmental structures to form during adult development. These AP-paired fibres that eventually define segmental boundaries show no correspondence to the orthogonal pattern of the nervous system until segments mature, when the transverse elements of both systems begin to register (indeed, expression of *tpm1* strikingly showed that myocyton populations in maturing segments become arranged in rows that correspond to the three transverse nerve cords in each segment; Fig. 2B). Being AP-paired, these muscle fibres provide a means to produce polarised expression of PCGs at segmental boundaries and it is likely that their associated myocytons account for some of the segmentally repeated PCG expression domains.

Additional repeated muscular expression domains are the circumferential foci that correspond to the outer and inner muscle layers, the central, medullary region where proglottids form, and the DV midline at intersegmental boundaries. The *sfl*+/*wnt*1+ signalling quartet and medial foci are also domains that could act as signalling centres associated with the onset of strobilation. Together with the marginal myocytons, these domains encompass all of the observed patterns of PCG expression, save expression at the nerve junctions seen in *hedgehog*, *sfl*, and *notum*.

In planarians it has been demonstrated that different muscle layers play distinct roles in Wnt signalling [15] and we can speculate that the different domains of expression, often relating spatially to different muscle layers, could organise different regions of the body. For example, the transverse fibres and/or signalling quartet could affect the organisation of the medullary region (i.e. proglottisation), whereas the cortical expression domains could be responsible for regionalising the AP axis of the body wall (i.e. strobilation). Similarly, expression of *sfrp* in the outer musculature of the neck could have a role in preventing ectopic initiation of segmentation, maintaining an unsegmented, generative region. Interestingly, Rozario et al. [40] demonstrated that anterior fragments of the neck region of *H. diminuta* have the ability to generate new proglottids in vitro, whereas very few were generated by middle and posterior fragments. Taking into account the results here, it is possible that in the posterior neck fragments only the transition zone that had already initiated segmentation was present, or that an unsegmented germinative region could not be stably maintained without signals elicited from the anterior of the neck.

Spatial domains of gene expression in tapeworms have only recently begun to be investigated [5,25,31,40] and specific markers for cell types are mostly lacking in this and other parasitic flatworm groups [31]. Nevertheless, the signalling domains identified here, such as the prominent pattern of the marginal myocytons, help to explain some previously published gene expression patterns [25,40].

### Wnt expression defines secondary, segmental axes in agreement with the polarity of the primary body axis

As in other animals, the components of Wnt signalling are expressed along the AP axis of *H*. *microstoma* in a highly regionalised pattern (Fig. 11) with the scolex and neck characterized by expression of the Wnt inhibitors, the transition zone by up-regulation of Wnt ligands, and the strobila by AP-polarised expression of ligands and inhibitors. These regions also represent developmental periodicities that illustrate the dynamic changes in signalling, with *wnt11a* being an example of a gene an especially limited periodicity of expression. Thus, the neck as classically defined is recognisable as a Wnt-inhibitory region, in which several frizzled receptors are expressed in staggered domains and in which a transition zone can be identified by the expression domain of *wnt11a*, by new, segmental domains of *sfl*, and by the up-regulation of *wnt1* and beginning of AP-polarised Wnt expression at segmental boundaries.

**Figure 11.**
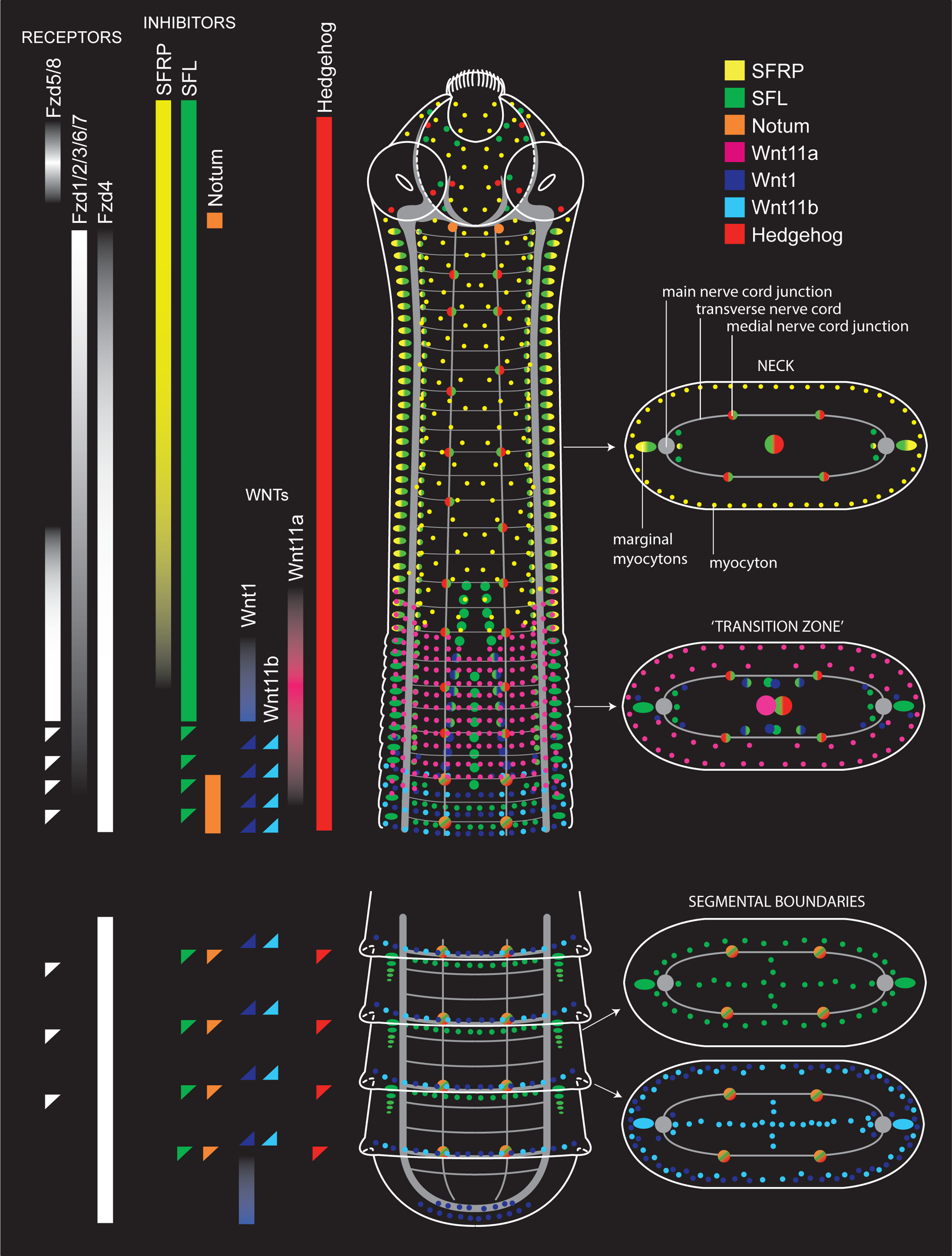
Diagram of Wnt and Hedgehog expression domains. Bars show regional expression domains along the AP axis while individual foci are shown in the illustration and cross sections. For clarity, the central medullary domains of expression in the neck region are depicted only in the cross sections. Multi-coloured foci depict co-localisation of the factors.

Previous investigation of Wnt signalling during larval development resolved historical questions regarding the polarity of the main body axis in tapeworms [6], demonstrating that the scolex is anterior as conventionally understood [5]. The question of the polarity of their individual segments in relation to the whole worm [7,8] is resolved by the present work which shows segmental boundaries marked by polarised expression of PCGs in agreement with the main body axis. The polarity of the tapeworm head-tail axis is therefore not dissimilar to other animals despite reduced cephalisation and unique features that confound anatomical comparisons.

### The ‘transition zone’ exhibits an embryological character

In planarians, orthologs of *wnt1* and *sfrp* have been reported to be co-expressed in cells during the initial stages of embryogenesis as well as during the earliest phases of regeneration, in which *sfrp* is expressed prior to being co-expressed with *wnt1* [13,43]. Co-localisation of *wnt1* and the inhibitor *notum* has also been demonstrated during the initial stages of regeneration, prior to their spatial domains becoming AP-polarised [44]. In *H. microstoma*, co-localisation of *wnt1* and *sfl* in the signalling quartet shows that Wnt signalling in the transition zone exhibits a character typical of early embryological and regenerative development in flatworms. Morphologically, the onset of strobilation is visible as a field of nascent segments (e.g. Fig. 5A) and we have never observed pre-strobilar worms with only a few or a single segment, indicating that strobilation acts across a field of tissues. The expression pattern of *wnt11a* corresponds to such a gradient and could act as a trigger to initiate strobilation. Moreover, being one of two *wnt11* paralogs present in flatworms [18] it may have been free to evolve a novel role in segmented tapeworms.

### Frizzled expression indicates Wnt signalling between muscle cells

Regional correspondence between ligands and antagonists and some Frizzled receptors suggests that specific binding partners may act to define different regions along the AP axis. In planarians, Scimone et al. [15] extended the work of Witchley et al. [14] to the different muscle layers showing that they play distinct roles during regeneration, with reduction in circular muscles causing medial-lateral axial defects and reduction in longitudinal muscles preventing re-establishment of positional information and failure to initiate regeneration. Their work identified a Wnt11 ligand and fzd5/8 receptor separately expressed by different subsets of musculature, suggesting signalling between muscle layers. Although we cannot definitively assign the expression of Wnt components to the different muscle layers here, concordance in the spatial expression patterns between *sfl* (Fig. 4) and *fzd5/8* (Fig. 7) points to signalling between muscle cells within and between different layers of their musculature.

### Neuronal expression of Hedgehog indicates a link with muscular Wnt signalling

Hedgehog is a secreted signalling protein in bilaterians that is typically expressed in gradients, with receiving cells having a concentration-dependent response [45,46]. It has pleiotropic effects during embryogenesis and is a key player in coordinating the development of the nervous system. In planarians, Hedgehog is expressed by differentiated neural cells in the ventral nerve cords and has been shown to have a role in establishing AP polarity by up-regulating Wnt signalling [29,47]. Similarly, expression of *hedgehog* in *H. microstoma* is observed in the nervous system, albeit in discrete foci in the cerebral ganglia and at the nerve junctions, as well as in the central region of the worm (Fig. 8). Although we cannot determine the identity of *hedgehog*+ cells in the central region, the foci could represent nerve cells or their progenitors associated with innervation of the proglottids.

Yazawa et al. [29] proposed that Hedgehog proteins in the CNS could be transported posteriorly via axonal trafficking [48] where they are secreted, stimulating Wnt expression in differentiated cells (e.g. myocytes) that in turn signal to neighbouring cells, causing up-regulation of β-catenin and a feedback loop that results in posteriorisation. *Wnt* and *hedgehog* expression in tapeworms is broadly consistent with this model, potentially linking Hedgehog signalling by the nervous system with Wnt expression by the musculature. Moreover, a number of different axonal vesicles have been described previously in the nervous system of *H. microstoma* [49] and exocrine secretion via nerve cells was recently demonstrated in diphyllobothriid tapeworms [50]. The main elements of the PNS also lie in intimate proximity to both the inner musculature and the layer of neoblast-like germinative cells (i.e. somatic stem cells) that are positioned at the cortical/medullary boundary [4], suggesting that as in planarians, these cells could be directed by signals elicited from the neuromuscular system.

During larval development *hedgehog* was found to be expressed along the dorsal and ventral midlines, perpendicular to the expression domain of *sfl* that mark the left and right axes (Figs. 8C-D). Midline expression is a conserved feature of Hedgehog signalling during embryogenesis that relates to CNS development [41,51] and involvement of Hedgehog in planarian neurogenesis has also been demonstrated [52,53]. As with Wnt expression [5], canonical expression of *hedgehog* during larval metamorphosis is consistent with this representing the phylotypic stage in tapeworm ontogeny [54].

### Tapeworm segmentation as a modified form of posterior regeneration in flatworms

Varying abilities of posterior regeneration is a widespread feature in flatworms [55] as all members of the group rely on the maintenance of pluripotent somatic stem cells for all cellular turnover [56]. The transition from larval to adult form in tapeworms can be compared most directly to planarian development by considering the case of posterior regeneration, as in both organisms this involves the loss and replacement of former posterior tissues. In planarians, loss of the posterior pole through amputation or fission [57] results in closure of the wound followed by the formation of a blastema in which new tissues are intercalated. Agata et al. [58] argued that the blastema was not simply a region containing undifferentiated (stem) cells, but rather that the initial development of the blastema re-established the lost (posterior) signalling centre which then induced regeneration of the missing tissues through a process of intercalation between the blastema and pre-existing anterior tissues, and grafting experiments showed that new tissues could be intercalated at the boundaries of the AP, DV and LR axes [58]. Almuedo-Castillo et al. [13] extended the model to include homeostasis, or ‘continuous intercalary re-specification’, to accommodate the fact that axial patterning systems such as Wnt are perpetually active in flatworms, even in fully intact, adult animals. In the transition to adult development, larval tapeworms lose their posterior cyst tissues in an analogous way to loss via amputation or fission and are left with only a rudimentary body prior to the development of the neck and strobila (Fig. 1A). We suggest the new posterior pole of the juvenile acts in a way similar to a blastema by becoming the new organiser. Consistent with this, during metamorphosis of the fox tapeworm, *Echinococcus multilocularis*, *wnt1* is expressed at the posterior pole of developing protoscoleces [5]. Post excystment, the neck and strobila are then continuously intercalated between the scolex and terminal tissues. We suggest that strobilar development in tapeworms is therefore also best understood as a type of continuous intercalary re-specification, differing in that an oscillating mechanism has been incorporated into the neck region to produce repetition of parts.

### Exadapting a gene regulatory network for segmentation

Historically, the morphogens Wnt1 and Hedgehog and the homeobox transcription factor Engrailed have been canonised as ‘segment polarity genes’ [10,59]. Conservation of Wnt-Hedgehog interactions in establishing boundaries between adjacent cells is a common mechanism in animals and has been used to infer a common origin of segmentation in bilaterians, or at least in protostomes [45,60], despite that the underlying modes of segmentation differ within and among different groups [10]. Tapeworm segmentation has been considered an evolutionary novelty [10], implying a lack of underlying homology to the GRNs of other animals. Planarians and other flatworms, as well as early-branching tapeworm lineages, lack segmentation. Nevertheless, AP patterning during planarian regeneration is dependent on Hedgehog-Wnt interactions [29,47] with knockdown phenotypes of Hedgehog mirroring the AP patterning defects of Wnt/β-catenin [61,62]. Despite their lack of segmentation, the interplay of Wnt and Hedgehog in planarian AP patterning has been considered homologous to that of segmented animals [45]. In tapeworms, Wnt and Hedgehog expression indicates that they also have a role in establishing and maintaining repeated AP axes along the strobila in addition to establishing their primary AP axis. Their involvement in polarising AP boundaries in tapeworms is thus perhaps unexpectedly similar to that of other protostomes, suggesting an underlying homology of their GRN or of specific regulatory modules used by bilaterians to produce boundaries between repeated parts.

The homeobox transcription factor Engrailed has been canonised as a ‘segmentation gene’ perhaps more strongly than any other due to the its role in forming parasegmental boundaries in *Drosophila* embryos, and although its function is mostly conserved in arthropods, it varies in other groups suggesting that a role in segmentation is not in fact ancestral [63]. An *H. microstoma* gene model containing an *engrailed*-like homeobox sequence (HmN_003004120) was first identified in the now complete assembly of its genome [28] and orthologous gene models are found in other parasitic and free-living flatworms. However, RNA-seq quantification in different ontogenetic stages and regions of the *H. microstoma* adult [25,26] demonstrate that *hm-engrailed* is either not expressed or expressed at levels below minimum thresholds (median 0-1.2 transcripts/million reads) throughout its ontogeny. Thus, although an *engrailed* ortholog is present in *H. microstoma* and broad presence of orthologs across the phylum point to its conservation, lack of expression makes it unlikely to be involved in tapeworm segmentation.

### Wnt signalling in relation to the evolution of segmentation and indeterminate growth in tapeworms

Segmentation is the hallmark of tapeworms but not a universally shared feature of the group, nor one likely to have been present in either their common ancestor or the earliest branching lineages of the clade [2,64]. True tapeworms (Eucestoda) are united by a universally observed larval form (the oncosphere) and other shared, derived characters irrespective of their final bauplan. The diversity of extant forms and the phylogenetic pattern of their evolution shows that the segmented body typical of the majority of contemporary groups is the result of two evolutionarily and developmentally independent processes: repetition of the hermaphroditic sets of reproductive organs (proglottisation) and their somatic compartmentalisation into semi-discrete segments (strobilation). Although coupled in most groups, there exist forms that demonstrate that tapeworms can be 1) neither proglottised nor segmented (e.g. caryophyllideans); 2) proglottised without being segmented (e.g. spathebothriideans); 3) lose segmentation secondarily without accompanying loss of proglottisation (e.g. *Anantrum* [64,65]); and 4) segmented without being proglottised (e.g. *Haplobothrium* [66]). These conditions demonstrate the independence of the processes, including the ability for them to be secondarily de-coupled, and phylogenetic analyses point to a step-wise pattern of evolution giving rise to the fully segmented condition found in most extant groups [2,64].

Overlaying strobilation on top of proglottisation more significantly allowed for the evolution of indeterminate growth, conferring fully segmented cestodes with significantly increased egg production and at least the theoretical ability for perpetual growth. Whereas spathebothriidean tapeworms evolved (or inherited) the ability to increase egg production through the development of multiple sets of reproductive organs (i.e. proglottisation), the number of proglottids they develop is determinate and ultimately restricted by their body size [67], which is in turn restricted by their host. In relation to fully segmented groups, their adult form can be viewed as an elongated neck region that fails to strobilate. The ability to compartmentalise the reproductive organs into segments that can then be progressively shed from the terminal end of the worm allowed for proglottisation to proceed indeterminately whilst their overall body size remained constant.

The existence of these forms considered in light of the present findings suggests a number of questions that merit further investigation. For example, the transverse muscle fibres that define segmental boundaries would not be expected to be observed in tapeworms that lack segmentation and/or proglottisation. Similarly, the clustered arrangement of the marginal myocytons could be a character restricted to groups that exhibit repetition of parts, or alternatively, that the presence or absence of PCG signalling, rather than the presence or absence of these elements of the musculature, could account for the differences in forms. The *wnt11a* paralog whose expression is associated with the onset of strobilation would be expected to play a different role in groups that lack this feature, and the presence of Wnt11 paralogs in flatworms [18] may have provided the opportunity for divergence in function. This work provides a PCG-based framework for addressing these questions through comparative investigations and the development of genomic resources representing most of the exemplar groups discussed above is currently underway.

## Conclusions

The unique combination of the maintenance of somatic stem cells and continuous expression of PCGs throughout life endows flatworms with extraordinary developmental plasticity, including partial and whole-body regeneration and highly complex life cycles and proliferative growth. Segmentation in tapeworms is a derived form of regeneration [40] that has been modified to produce repetition of the reproductive organs. PCG expression during strobilar growth shows that segmental boundaries are marked by polarised expression of conserved patterning genes used throughout the animal kingdom to produce repeated body parts. Similarly, the AP polarity of their bodies and of their individual segments is found to be the same as that of other animals. These finding suggest that despite extreme differences in the trophic and life history strategies of parasitic flatworms, the GRNs that pattern their bodies do not differ fundamentally from their free-living cousins either within or outside of the phylum. Thus the study of parasitic species can be instructive not only for understanding unique aspects of their own biology and the diseases they cause, but of animals more generally. For example, genomic investigations have shown that stem cell pluripotency in parasitic flatworms is maintained in the absence of putatively essential, ‘universal’ multipotency proteins such as Piwi, Vasa [68,69] and Wnt3 [18,70], while complete assembly of the *H. microstoma* genome demonstrated that their chromosomes are capped by centromeric motifs, making them responsible not only for chromosome segregation but also for the protection of ends normally conferred by telomeres [28]. Such unexpected findings speak to general principles in biology and not to the specific features of the groups investigated or the diseases they cause, illustrating the value of broadening the range of model organisms that shape our understanding of biology [3,71,72].

## Methods

### Animals

Adult and larval specimens of the Nottingham strain [20] of *Hymenolepis microstoma* were generated, fixed and stored as described in [25]. Mice were used in accordance with animal care regulations under U.K. Home Office license PPL70/8684.

### Genes and probe synthesis

Gene orthologs were identified from *H. microstoma* genomic data and gene models as previously described [5,18,21]. An isoform of *collagen* was chosen as it was orthologous to the isoform investigated in the planarian *Schmidtea mediterranea* (Smed_00066_V2; [14]). A list of the gene models, primers and protein sequences investigated here is given in Additional file 1. Primers were designed against the gene models using Primer3 [73] implemented in Geneious (Biomatters Ltd) and near full-length mRNA transcripts amplified by PCR from cDNAs synthesised from total RNA purified from either larval or adult samples. Amplicons were cloned into StrataClone (Agilent Technologies) or pGEM-T (Promega) vectors and eight positive colonies/gene transferred to 750 ul of ddH2O and heated to 90 C to liberate the plasmids. The size and direction of the inserts were then checked via PCR by combining M13R with GSP forward and reverse primers, and the identities of the inserts confirmed via Sanger sequencing. Insert regions together with flanking T3/T7 or T7/SP6 promotor sites were then amplified in large volume (75 ul) reactions using M13F/R or T7/SP6 primers. Resulting amplicons were used as templates for the synthesis of either digoxigenin (DIG) or fluorescein (FITC)-labelled anti-sense probes by *in vitro* transcription using T7 or T3 polymerases (Roche) and DIG or FITC RNA labelling mixes (Roche).

### Fluorescent *in situ* hybridisation (FISH)

Tyramide-fluorescein-based FISH was performed with digoxigenin-labelled antisense probes as previously described [5]. Double FISH was performed using a modified version of this procedure. During probe hybridisation, both DIG and FITC-labelled antisense probes (each at a concentration of 1 µg/mL) were hybridised simultaneously. FITC-labelled probes were detected first by incubating with sheep Anti-Fluorescein-POD Fab fragments (Roche) at a 1:50 dilution. The signal was then developed via tyramide signal amplification (TSA) using a fluorescein tyramide solution as previously described [5] followed by incubation in 100 mM sodium azide for 45 min to quench the HRP (horseradish peroxidase) enzyme. Samples were then washed four times in PBST (phosphate buffered saline with 1% Tween) for 10 min before incubating in blocking buffer for one hour. Digoxigenin-labelled probes were then detected by incubating with sheep Anti-Digoxigenin-POD Fab fragments (Roche) at a 1:50 dilution. The signal was then developed via TSA using a rhodamine tyramide solution. Specimens were then counterstained in DAPI (4’,6-diamidino-2-phenyllindole), cleared in 80% glycerol and mounted on microscope slides.

### Immunohistochemistry

Phalloidin staining was done on PFA-fixed specimens rinsed in PBS and permeabilised using PBS containing 0.25% TritonX-100 for 2h. Specimens were stained with AlexaFluor-488 labelled phalloidin (Molecular Probes) in PBS at a final concentration of 1U/ml. After treatment with Ribonuclease A to remove RNA, nuclear counterstaining was carried out with propidium iodide (1:250) for 20 min. Specimens were rinsed again and mounted on glass slides in 90% glycerol, 10% PB, 0.25% DABCO.

Staining against Synapsin was combined with TSA and performed on PFA-fixed specimens permeabilised in 1% sodium dodecyl sulfate (N.B. no Proteinase K treatment) for one hour prior to the FISH procedure. After FISH detection, specimens were quenched in 100 mM sodium azide to prevent cross-reactivity of the tyramide solutions. Anti-synapsin (3C11 anti-SYNORF1, Developmental Studies Hybridoma Bank) was used as the primary antibody at a 1:200 dilution. The second antibody was goat-anti-mouse conjugated to HRP. The signal was then amplified via TSA with a rhodamine tyramide solution, as described in the FISH procedure. DAPI (Thermofisher) counterstaining was performed last on all specimens by incubating in a 4 ng/mL solution for 10 min.

### Imaging

Brightfield and fluorescent imaging was done using a Leica DM5000B epifluorescent microscope with CoolLED illumination and a DFC450C digital camera linked to Leica Application Suite ver. 4. Confocal imaging was performed with a Nikon A1-Si confocal microscope and maximum projections and 3D reconstructions made from resulting image stacks using the Fiji distribution of ImageJ2 [74,75]. In some cases increased differentiation of signals was achieved by adjusting the overall brightness of different channels.

### Additional files

**Additional file 1.** Table of gene models and primers.

**Additional file 2.** Movie: 3D projection of *collagen* expression in Stage 3 larva.

**Additional file 3.** Movie: 3D projection of *collagen* expression in encysted, Stage 4 larva.

**Additional file 4.** Movie: 3D projection of *sfl* expression in the transition zone.

**Additional file 5.** Movie: 3D projection of *sfl* (green) + *wnt1* (magenta) expression in the transition zone.

**Additional file 6.** Movie: 3D projection of *sfl* (green) + *wnt1* (red) expression in the strobila.

**Additional file 7.** Movie: 3D projection of *wnt11a* expression in the transition zone.

**Additional file 8.** Movie: 3D apical projection of *hedgehog* (magenta) + *sfl* (green) expression in Stage 1 larva.

**Additional file 9.** Movie: 3D projection of *hedgehog* expression in Stage 1 larva.

**Additional file 10.** Movie: 3D projection of *hedgehog* expression in Stage 3 larva.

**Additional file 11.** Movie: 3D projection of *hedgehog* expression in encysted, Stage 4 larva.

## Availability of data and materials

All genome data are available via WormBase ParaSite (http://parasite.wormbase.org) [76].

## Author contributions

PDO and FJ designed the study with input from UK. FJ led and conducted the study. NR performed initial Wnt identification, cloning and colorimetric WMISH. FJ, AB, JM and UK generated FISH results. PDO, UK and FJ interpreted the results. PDO prepared and wrote the paper with input from UK. All authors read and approved the submitted version.

## Acknowledgements

We are grateful to Maximilian Telford for providing co-supervisory support to FJ. We thank Alex Gruhl for phalloidin staining and imaging, Maria Abellas-Noguerol for anti-Synapsin staining and imaging and Edward O’Garro-Priddie for generating initial FISH results for *collagen*. The anti-Synapsin hybridoma developed by E. Buchner, University of Würzburg, Germany, was obtained from the Developmental Studies Hybridoma Bank, created by the NICHD of the NIH and maintained at The University of Iowa, Department of Biology, Iowa City, IA 52242.

